# Genetic loci associated with prevalent and incident myocardial infarction and coronary heart disease in the Cohorts for Heart and Aging Research in Genomic Epidemiology (CHARGE) Consortium

**DOI:** 10.1101/2020.02.21.959312

**Authors:** Julie Hahn, Yi-Ping Fu, Michael R. Brown, Joshua C. Bis, Paul S. de Vries, Mary F. Feitosa, Lisa R. Yanek, Stefan Weiss, Franco Giulianini, Albert Vernon Smith, Xiuqing Guo, Traci M. Bartz, Diane M. Becker, Lewis C. Becker, Eric Boerwinkle, Jennifer A. Brody, Yii-Der Ida Chen, Oscar H. Franco, Megan Grove, Tamara B. Harris, Albert Hofman, Shih-Jen Hwang, Brian G. Kral, Lenore J. Launer, Marcello R. P. Markus, Kenneth M. Rice, Stephen S. Rich, Paul M. Ridker, Fernando Rivadeneira, Jerome I. Rotter, Nona Sotoodehnia, Kent D. Taylor, André G. Uitterlinden, Uwe Völker, Henry Völzke, Jie Yao, Daniel I. Chasman, Marcus Dörr, Vilmundur Gudnason, Rasika A. Mathias, Wendy Post, Bruce M. Psaty, Abbas Dehghan, Christopher J. O’Donnell, Alanna C. Morrison

**Affiliations:** Human Genetics Center, Department of Epidemiology, Human Genetics, and Environmental Sciences, School of Public Health, The University of Texas Health Science Center at Houston, Houston, Texas USA; Office of Biostatistics Research, National Heart, Lung, and Blood Institute, National Institutes of Health, Bethesda, Maryland, USA; Framingham Heart Study, National Heart, Lung, and Blood Institute, National Institutes of Health, Framingham, Massachusetts, USA; Cardiovascular Health Research Unit, Department of Medicine, University of Washington, Seattle, Washington, USA; Department of Epidemiology, Erasmus University Medical Center, Rotterdam, The Netherlands; Division of Statistical Genomics, Department of Genetics, Washington University School of Medicine, St. Louis, Missouri, USA; GeneSTAR Research Program, Department of Medicine, Johns Hopkins University School of Medicine, Baltimore, Maryland, USA; Interfaculty Institute for Genetics and Functional Genomics, The University Medicine and Ernst-Moritz-Arndt-University Greifswald, Greifswald, Germany; DZHK (German Centre for Cardiovascular Research), partner site Greifswald, Greifswald, Germany; Division of Preventive Medicine, Brigham and Women’s Hospital, Boston, Massachusetts, USA; Icelandic Heart Association, Kovapvogur, Iceland; Faculty of Medicine, University of Iceland, Reykajvik, Iceland; The Institute for Translational Genomics and Population Sciences, Department of Pediatrics, The Lundquist Institute for Biomedical Innovation at Harbor-UCLA Medical Center, Torrance, California, USA; Department of Biostatistics, The University of Washington, Seattle, Washington, USA; Human Genome Sequencing Center, Baylor College of Medicine, Houston, Texas, USA; Department of Internal Medicine, Erasmus University Medical Center, Rotterdam, The Netherlands; Laboratory of Epidemiology and Population Sciences, Intramural Research Program, National Institute on Aging, National Institutes of Health, Bethesda, Maryland, USA; Department of Epidemiology and Biostatistics, Imperial College London, London, United Kingdom; Department of Internal Medicine B - Cardiology, Pneumology, Infectious Diseases, Intensive Care Medicine, The University Medicine Greifswald, Greifswald, Germany; Department of Medicine and Epidemiology, Johns Hopkins University School of Medicine, Baltimore, Maryland, USA; Harvard Medical School, Boston, Massachusetts, USA; Division of Cardiology, Department of Medicine, University of Washington, Seattle, Washington, USA; Department of Epidemiology, Harvard T.H. Chan School of Public Health, Boston, Massachusetts, USA; Institute for Community Medicine, University Medicine Greifswald, Greifswald, Germany; Department of Epidemiology, The University of Washington, Seattle, Washington, USA; Department of Health Services, The University of Washington, Seattle, Washington, USA; Kaiser Permanente Research Institute, Seattle, Washington, USA; VA Boston Healthcare System, Veteran’s Affair, Boston, Massachusetts, USA; Cardiovascular Medicine Division, Brigham and Women’s Hospital, Boston, Massachusetts, USA

## Abstract

**Background:** Genome-wide association studies have identified multiple genomic loci associated with coronary artery disease, but most are common variants in non-coding regions that provide limited information on causal genes and etiology of the disease. To better understand etiological pathways that might lead to discovery of new treatments or prevention strategies, we focused our investigation on low-frequency and rare sequence variations primarily residing in coding regions of the genome while also exploring associations with common variants.

**Methods and Results:** Using samples of individuals of European ancestry from ten cohorts within the Cohorts for Heart and Aging Research in Genomic Epidemiology (CHARGE) consortium, both cross-sectional and prospective analyses were conducted to examine associations between genetic variants and myocardial infarction (MI), coronary heart disease (CHD), and all-cause mortality following these events. Single variant and gene-based analyses were performed separately in each cohort and then meta-analyzed for each outcome. A low-frequency intronic variant (rs988583) in *PLCL1* was significantly associated with prevalent MI (OR=1.80, 95% confidence interval: 1.43, 2.27; *P*=7.12 × 10^-7^). Three common variants, rs9349379 in *PHACTR1*, and rs1333048 and rs4977574 in the 9p21 region, were significantly associated with prevalent CHD. Four common variants (rs4977574, rs10757278, rs1333049, and rs1333048) within the 9p21 locus were significantly associated with incident MI. We conducted gene-based burden tests for genes with a cumulative minor allele count (cMAC) ≥ 5 and variants with minor allele frequency (MAF) < 5%. *TMPRSS5* and *LDLRAD1* were significantly associated with prevalent MI and CHD, respectively, and *RC3H2* and *ANGPTL4* were significantly associated with incident MI and CHD, respectively. No loci were significantly associated with all-cause mortality following a MI or CHD event.

**Conclusion:** This study confirmed previously reported loci influencing heart disease risk, and one single variant and three genes associated with MI and CHD were newly identified and warrant future investigation.

## Introduction

Coronary heart disease (CHD) is a leading cause of morbidity and mortality worldwide, accounting for one of every seven deaths in the United States in 2016. (1) In addition to major modifiable risk factors such as dyslipidemia, hypertension, diabetes, and cigarette smoking (2), genetic susceptibility to CHD has also been investigated extensively through family-based studies, candidate gene studies, and more recently genome-wide association studies (GWAS). (3-9) With progressively expanded sample sizes in recent GWAS, at least 160 loci have been associated with the risk of coronary artery disease. (10-13) Most of these loci are represented by common variants located in noncoding regions, resulting in limited implications for causal genes and etiological pathways. Further, while most available data are derived from genome-wide analysis of prevalent CHD, data are sparse from prospective studies of incident cardiovascular events in populations free of baseline cardiovascular disease.

Low-frequency and rare coding sequence variations across the genome have been investigated in studies of cardiovascular disease risk factors (14-18), with the goal of better understanding the etiology of these risk factors and to advance the discovery of the treatment and prevention of diseases. (19) We previously published the results from a prospective analysis of CHD among individuals of European ancestry from the Cohorts for Heart and Aging Research in Genomic Epidemiology (CHARGE) Consortium, and identified low-frequency and common variants associated with incident CHD. (20)

In this current study of individuals of European ancestry, we implemented both a cross-sectional and prospective study design in the setting of the CHARGE Consortium to examine the association between genetic variants and the risk of prevalent and incident myocardial infarction (MI) and CHD. Study of incident cardiovascular events is enabled by the rigorous prospective design of population cohorts contributing to the CHARGE Consortium. We also investigated whether these genetic variants are associated with all-cause mortality after incident MI and CHD.

## Materials and Methods

### Study design and participants

Ten cohorts within the CHARGE Consortium Subclinical Working Group were included in this study: Age, Gene, Environment, Susceptibility Study (AGES), Atherosclerosis Risk in Communities (ARIC) Study, Cardiovascular Health Study (CHS), Family Heart Study (FamHS), Framingham Heart Study (FHS), the GeneSTAR Study (GeneSTAR), Multi-Ethnic Study of Atherosclerosis (MESA), Rotterdam Study (RS), Study of Health in Pomerania (SHIP), and the Women’s Genome Health Study (WGHS). Detailed characteristics of the participating cohorts and study participant are shown in the **Supporting Document.** All study participants provided written informed consent to participate in genetic studies, and all study sites received approval to conduct this research from their local Institutional Review Boards (IRB) respectively.

### Genotype calling and quality control

Participants from WGHS were genotyped by the HumanHap300 Duo+ (Illumina, Inc., San Diego, CA), and all other study participants were genotyped by the HumanExome BeadChip (v1.0-1.2, Illumina, Inc., San Diego, CA) which contains more than 240,000 variants including those discovered through exome sequencing in ∼12,000 individuals and other non-coding common variants such as previously-reported GWAS signals and ancestry-informative markers. Data for AGES, ARIC, CHS, FamHS, FHS, MESA, and RS were jointly called at the University of Texas Health Science Center at Houston (21); SHIP was called in Illumina GenomeStudio using the CHARGE Consortium joint calling cluster file; GeneSTAR used the Illumina GenomeStudio and zCall software (22); and WGHS data was called using the Illumina BeadStudio v.3.3. Variant quality control (QC) was performed centrally (21) and by the individual studies, including checking concordance with previous GWAS data, and excluding participants with missing >5% genotypes, population clustering outliers, individuals with high inbreeding coefficients or heterozygote rates, gender mismatches, duplicated pairs, and unexpectedly high proportion of identity-by-descent sharing for family studies.

### Cardiovascular outcome definition

Two cardiovascular outcomes were examined for association in this study: 1) MI: fatal or non-fatal MI; and 2) CHD: fatal or non-fatal MI, fatal CHD, sudden death within one hour of onset of symptoms, or revascularization (percutaneous coronary artery intervention such as stent or balloon angioplasty, or coronary artery bypass grafting). No exclusions were applied for the cross-sectional analysis of prevalent MI and prevalent CHD. For analysis of incident events, participants with a history of MI, CHD or revascularization at the baseline examination were excluded. All-cause mortality after MI or CHD was also investigated with follow-up time from first MI or CHD incident events until death, loss to follow-up, or the end of study.

### Statistical analysis

Single variant and gene-based analyses were conducted in each participating cohort respectively, followed by meta-analysis performed for each cardiovascular outcome to summarize results. All autosomal variants were coded to the minor allele observed in the CHARGE jointly called data (21) and assumed log-additive genetic effect in the analyses. The minor allele frequency (MAF) thresholds were defined using the European allele frequencies derived from the CHARGE jointly called data. (21) Variant annotation was performed centrally within CHARGE using dbNSFP. (23, 24) Variants with MAF ≥ 1% were included in single variant tests for prevalent MI and CHD and for incident MI. Single variant results for incident CHD followed the same analytic approach and are reported in Morrison et al. (20) and are not reported in detail here. Gene-based tests were evaluated for MI and CHD outcomes: the Sequence Kernel Association Test (SKAT) (25) and a burden test (26). Only functional coding variants (missense, stop-gain, stop-loss, or splice-site changes) with MAF < 5% were aggregated by gene, and we only analyzed genes with a cumulative minor allele count (cMAC) ≥ 5.

For both single variant and gene-based burden tests of prevalent events, we performed Firth’s logistic regression model to test the association between each variant and cardiovascular outcome using the “logistf” package in R (27-29) to account for the possible inflated type one error in the rare variant association analysis in a case-cohort study design.(30) Meta-analysis for prevalent events was conducted with METAL (31) and applied the genomic control correction. For the single variant and two gene-based tests of incident events, a Cox proportional hazards regression model implemented in the seqMeta package in R was used to test the association between each variant and the incident event or post-event all-cause mortality. SeqMeta was used both at the study-specific analysis and meta-analysis levels. (32) All study-specific analyses (single variant and gene-based tests) were adjusted for cohort-specific design variables (e.g. study sites, family structure) and for population substructure using principal components as needed. We applied a Bonferroni corrected threshold to determine statistical significance in each analysis as described below.

## Results

### Prevalent MI and CHD association

A total of 27,349 participants of European ancestry from seven cohorts including 1831 prevalent MI cases (6.7%) and 2518 prevalent CHD cases (9.2%) were used in the meta-analyses of prevalent events (**S1 Table**). We examined individually a total of 36,406 variants, combining both low-frequency and common variants (MAF ≥ 1%), across all autosomal chromosomes corresponding to a Bonferroni corrected significance threshold of *P*=1.37 × 10^-6^. A low-frequency (MAF=1.64%) intronic variant (rs988583) in the phospholipase C like 1 gene (*PLCL1*) was significantly associated with prevalent MI (*P*=7.12 × 10^-7^; OR=1.80, 95% confidence interval=1.43 to 2.27; **Table 1**). Three common variants were significantly associated with prevalent CHD: rs9349379 in *PHACTR1* and rs1333048 and rs4977574 in the 9p21 region (**Table 1**).

**Table 1.**
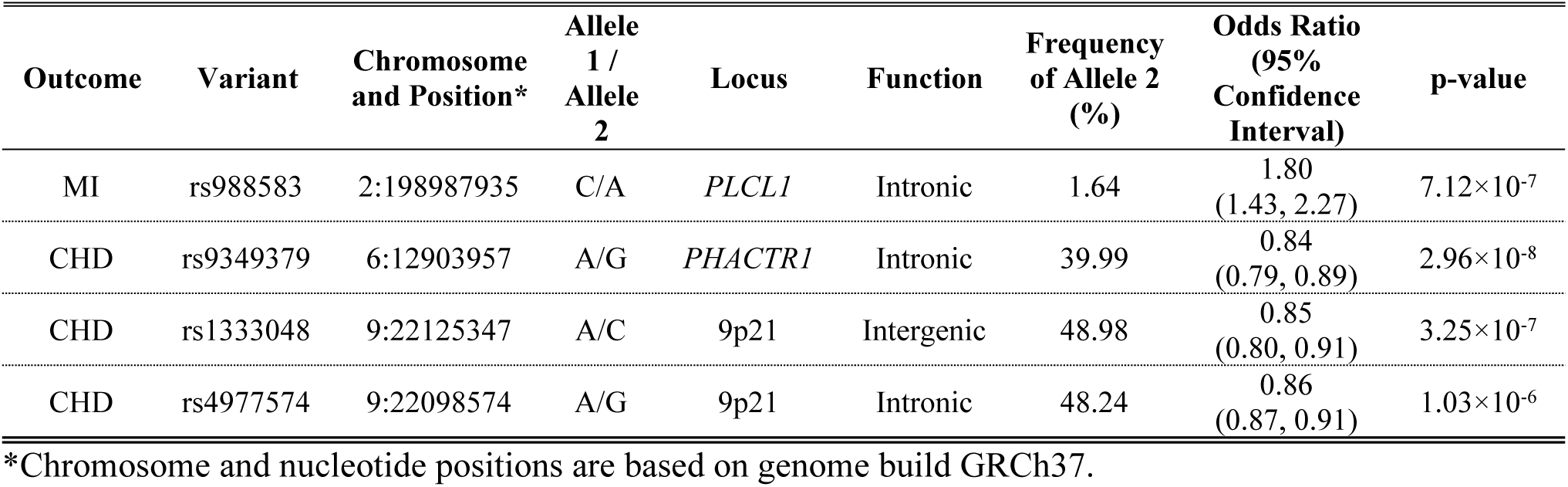
Low-frequency and common variants associated with prevalent MI and CHD.

| Outcome | Variant | Chromosome and Position* | Allele 1 / Allele 2 | Locus | Function | Frequency of Allele 2 (%) | Odds Ratio (95% Confidence Interval) | p-value |
| --- | --- | --- | --- | --- | --- | --- | --- | --- |
| MI | rs988583 | 2:198987935 | C/A | *PLCL1* | Intronic | 1.64 | 1.80<br>(1.43, 2.27) | 7.12×10<sup>-7</sup> |
| CHD | rs9349379 | 6:12903957 | A/G | *PHACTR1* | Intronic | 39.99 | 0.84<br>(0.79, 0.89) | 2.96×10<sup>-8</sup> |
| CHD | rs1333048 | 9:22125347 | A/C | 9p21 | Intergenic | 48.98 | 0.85<br>(0.80, 0.91) | 3.25×10<sup>-7</sup> |
| CHD | rs4977574 | 9:22098574 | A/G | 9p21 | Intronic | 48.24 | 0.86<br>(0.87, 0.91) | 1.03×10<sup>-6</sup> |

\*Chromosome and nucleotide positions are based on genome build GRCh37.

In the gene-based burden tests, we analyzed 16,628 autosomal genes that contained functional low-frequency or rare variants with MAF < 5% and with a cumulative minor allele count (cMAC) ≥ 5; therefore, the Bonferroni corrected p-value threshold was *P*=3.01 × 10^-6^. The transmembrane serine protease 5 gene (*TMPRSS5*) on chromosome 11, containing nine nonsynonymous rare variants (**S2 Table**), was significantly associated with prevalent MI (*P*=2.59 × 10^-6^, OR=3.00, 95% confidence interval: 1.90, 4.73; **Table 2**). The low-density lipoprotein receptor class A domain containing 1 gene (*LDLRAD1*) on chromosome 1 contained seven rare variants (**S2 Table**) and was significantly associated with prevalent CHD (*P*=1.30 × 10^-6^, OR=4.48, 95% confidence interval: 2.44, 8.23; **Table 2**).

**Table 2.** Genes associated with prevalent MI and CHD in gene-based analysis.

| Outcome | Gene | Chromosome and Position* | cMAC** | Variants (n)^ | Test | Odds Ratio (95% Confidence Interval) | p-value |
| --- | --- | --- | --- | --- | --- | --- | --- |
| MI | *TMPRSS5* | 11:113558268-113577151 | 152.02 | 9 | Burden | 3.00 (1.90, 4.73) | 2.59×10<sup>-6</sup> |
| CHD | *LDLRAD1* | 1:54472971-54483859 | 60.05 | 7 | Burden | 4.48 (2.44, 8.23) | 1.30×10<sup>-6</sup> |

\*Chromosome and nucleotide positions are based on genome build GRCh37.
\*\*cMAC = overall cumulative minor allele count.
^Variants (n) = number of variants included in the analysis; variants were restricted to those with MAF < 5% and annotated as nonsynonymous, splice-site, or stop loss/gain function.

### Incident MI and CHD association

Nine cohorts contributed a total of 55,736 participants of European ancestry to the analyses of incident events, where 3,031 incident MI cases (5.4%) were reported during an average of 15.0 years of follow-up and 5,425 incident CHD cases (9.73%) were reported during an average of 15.6 years of follow-up (**S3 Table**). A total of 37,109 low frequency and common autosomal variants (MAF ≥ 1%) were individually tested for association with incident MI, with adjustment of age, sex, and population substructure. The Bonferroni corrected p-value threshold for single variant analysis of incident MI was *P*=1.35 × 10^-6^. Four common variants in the noncoding region at the 9p21 locus were significantly associated with incident MI (**Table 3**). As previously stated, single variant results for incident CHD are reported in Morrison et al. (20) and are not reported here.

**Table 3.**
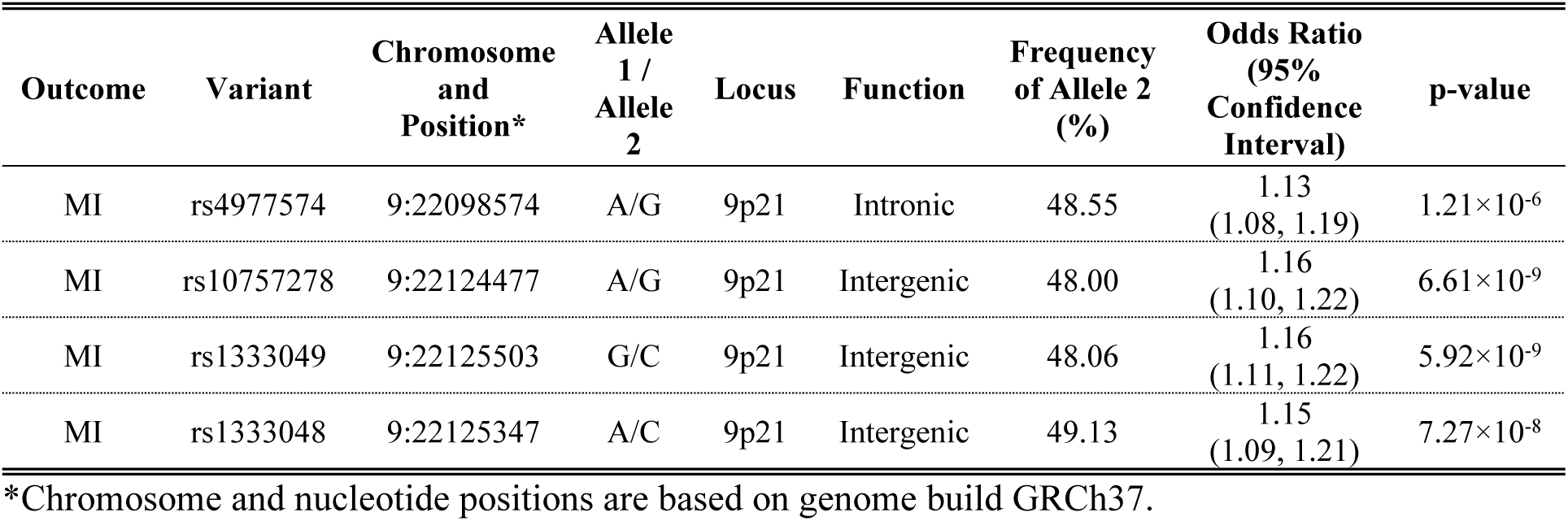
Low-frequency and common variants associated with incident MI.

| Outcome | Variant | Chromosome and Position* | Allele 1 / Allele 2 | Locus | Function | Frequency of Allele 2 (%) | Odds Ratio (95% Confidence Interval) | p-value |
| --- | --- | --- | --- | --- | --- | --- | --- | --- |
| MI | rs4977574 | 9:22098574 | A/G | 9p21 | Intronic | 48.55 | 1.13<br>(1.08, 1.19) | $1.21 \times 10^{-6}$ |
| MI | rs10757278 | 9:22124477 | A/G | 9p21 | Intergenic | 48.00 | 1.16<br>(1.10, 1.22) | $6.61 \times 10^{-9}$ |
| MI | rs1333049 | 9:22125503 | G/C | 9p21 | Intergenic | 48.06 | 1.16<br>(1.11, 1.22) | $5.92 \times 10^{-9}$ |
| MI | rs1333048 | 9:22125347 | A/C | 9p21 | Intergenic | 49.13 | 1.15<br>(1.09, 1.21) | $7.27 \times 10^{-8}$ |

\*Chromosome and nucleotide positions are based on genome build GRCh37.

For the gene-based analyses, we examined 17,574 genes across all autosomal chromosomes for association with incident MI, and the Bonferroni corrected significance level was *P*=2.85 × 10^-6^. The ring finger and CCCH-Type domains 2 gene (*RC3H2*) on chromosome 9 was significantly associated with incident MI in the burden test (*P*=2.99 × 10^-6^, OR=0.35, 95% confidence interval=0.23, 0.55; **Table 4**) and contained 12 nonsynonymous and one splice-site rare variants (**S4 Table**). No genes were significantly associated with incident MI using SKAT. For the gene-based analyses of incident CHD, 16,620 genes were evaluated and the Bonferroni significance levels was P=3.01 × 10^-6^. Angiopoietin-like 4 (*ANGPTL4*) on chromosome 19 was significantly associated with incident CHD using SKAT (P=1.29 × 10^-6^; **Table 4**) and contained 10 variants (**S4 Table**), and no gene was significantly associated using the burden test.

**Table 4.** Genes associated with incident MI and CHD in gene-based analysis.

| Outcome | Gene | Chromosome and Position* | cMAC** | Variants (n)^ | Test | Odds Ratio (95% Confidence Interval) | p-value |
| --- | --- | --- | --- | --- | --- | --- | --- |
| MI | *RC3H2* | 9:125606835-125667562 | 356.02 | 13 | Burden | 0.35 (0.23, 0.55) | $2.99 \times 10^{-6}$ |
| CHD | *ANGPTL4* | 19:8429011-843925 | 2830.07 | 10 | SKAT | - | $1.29 \times 10^{-6}$ |

\*Chromosome and nucleotide positions are based on genome build GRCh37.
\*\*cMAC = overall cumulative minor allele count.
^Variants (n) = number of variants included in the analysis; variants were restricted to those with MAF < 5% and annotated as nonsynonymous, splice-site, or stop loss/gain function.

### Post MI and CHD mortality analysis

Among the 3,751 MI and CHD cases from six cohorts that contributed to the analysis of all-cause mortality, there were 1,860 all-cause deaths over a mean 10.9 years of follow-up (**S5 Table**). We examined 36,685 autosomal variants with MAF ≥ 1% in the single variant analysis (Bonferroni corrected significant level of *P*=1.36 × 10^-6^) and 17,574 genes in the gene-based analysis (Bonferroni corrected significant level of *P*=2.85 × 10^-6^). No single variant or gene reached the significance threshold in the analysis of all-cause mortality among survivors of MI or CHD. We examined the significant variants and genes reported in Tables 1-4 for their relationship with mortality following a MI or CHD event (**S6 Table**). While these loci were significantly associated with prevalent and incident MI or CHD events, only the 9p21 common variants were nominally associated with all-cause mortality (P < 0.05). The 9p21 variants that were associated with reduced risk of prevalent CHD (rs1333048 and rs4977574; **Table 1**), and with increased risk of incident MI (rs1333048, rs4977574, rs10757278, and rs1333049; **Table 3**) were all associated with modestly reduced risk of all-cause mortality (**S6 Table**).

## Discussion

Our study evaluated genetic susceptibility to MI and CHD in cross-sectional and prospective settings among individuals of European ancestry. We confirmed several previously reported loci and newly identified one low-frequency variant and three genes harboring low-frequency and rare coding variants that warrant investigation in future studies.

Single variant analysis of prevalent cardiovascular outcomes revealed a low-frequency (MAF=1.64%) intronic variant, rs988583, in *PLCL1* significantly associated with increased risk of MI (*P*=7.12 × 10^-7^). *In silico* replication was conducted by a look up of rs988583 and its association with prevalent MI in the Myocardial Infarction Genetics and CARDIoGRAM exome chip meta-analysis public release (33), and there was no significant association with MI (*P*=0.34). A GWAS of MI and coronary artery disease (CAD) in a Saudi Arab population identified an intergenic variant, rs7421388, near *PLCL1* associated with CAD (*P* = 4.31 × 10^-6^) and replicated in an independent sample of Saudi Arabs (*P* = 5.37 × 10^-7^). (34) In another study of an ethnic Arab population, rs1147169 in *PLCL1* was protective against a low level of high density lipoprotein-cholesterol levels (*P* = 2.87 × 10^-7^). (35) In individuals of European ancestry, rs988583 and rs1147169 are in linkage equilibrium (R^2^= 0.0043). In addition to these studies, *PLCL1* has been implicated in coronary artery aneurysm in Kawasaki disease and *PLCL1* might play a role in the regulation of vascular endothelial cell inflammation via interference with proinflammatory cytokine expression. (36)

A burden test aggregating low-frequency and rare coding variants in genes showed a significant positive association between *TMPRSS5* and prevalent MI (*P*=2.59 × 10^-6^) and *LDLRAD1* and prevalent CHD (P=1.30 × 10^-6^), and a significant protective association between *RC3H2* and incident MI (*P*=2.99 × 10^-6^). A significant association between *ANGPTL4* and incident CHD was identified using SKAT (P=1.29 × 10^-6^). The relationship between *ANGPTL4* and CHD has been previously reported, with the rs116843064 missense variant playing a major role in reducing lipid levels and risk of CHD. (33, 37) Serine proteases, such as *TMPRSS5*, are known to be involved in many physiological and pathological processes, and *TMPRSS5* has been implicated in impaired hearing function. (38) Little is known about *LDLRAD1*, with most marked gene expression in lung and fallopian tube (39), and a rare variant in this gene has been associated with breast cancer. (40) Roquin-2 is encoded by *RC3H2* and has been shown to play a key role in posttranscriptional regulation of autoimmunity and inflammatory response. (41) Each of these genes associated with prevalent or incident cardiovascular outcomes has rare and low-frequency variants underlying the gene burden tests (**S2 and S4 Table**). We identified 11 putative driving variants of these gene-based associations (i.e. those with p<0.05 in **S2 and S4 Table**; rs201233178, rs200417674, and rs116913282 in *TMPRSS5*; rs150560713, rs202234131, rs142900519, and rs76122098 in *LDLRAD1*; rs201920127, rs144714368, and rs199901510 in *RC3H2*; and rs116843064 in *ANGPTL4*). An *in silico* replication was not possible due to the rare frequency of these coding variants and their absence in the public release of the Myocardial Infarction Genetics and CARDIoGRAM exome chip meta-analysis or the analysis of CAD in the UK Biobank and the UK Biobank and CARDIoGRAMplusC4D meta-analysis (10, 33) However, it is important to note that rs116843064 of *ANGPTL4* is the same variant found in the single variant analysis conducted for incident CHD by Morrison et al., and this gene is likely to be driving the significant association found in the SKAT analysis of incident CHD. (20) It is of interest that the effect sizes of the gene-based tests (**Tables 2 and 4**) are larger than the single variant test effect sizes (**Tables 1 and 3**), supporting the notion that low-frequency and rare variants may have a more substantial impact on disease risk.

Although there was no statistically significant result found for all-cause mortality after MI or CHD, after accounting for multiple testing, the protective direction of effect for many of our mortality results suggests that genetic variants might contribute differently in various stages of disease manifestation. Specifically, our results highlight differences in the direction of effect for common variants at the 9p21 locus associated with decreased risk of prevalent CHD, increased risk of incident MI, and a nominally significant reduced risk of all cause-mortality following a cardiovascular event. Generally, the loci identified for prevalent disease were not the same as those identified for incident disease, as has been observed in previous studies. (9) Indeed, comparison of prevalent and incident findings (**S6 Table**) shows that the single variant (*PLCL1* locus) and gene-based (*TMPRSS5* locus) results for prevalent MI were not significantly associated with incident MI, and the direction of effects were consistent for *PLCL1* but not *TMPRSS5*. Similarly, the significant gene, *LDLRAD1*, identified for increased risk of prevalent CHD was not significantly associated with incident CHD, but the direction of effect was consistent. The *RC3H2* gene, which showed an inverse association with incident MI, was not significantly associated with prevalent MI and it exhibited an opposite direction of effect. A possible explanation for these observed differences is that genetic studies of cardiovascular diseases are usually conducted with the cross-sectional study design, which has the potential to oversample participants with longer post-event survival (42) and the results do not always replicate in the prospective studies for disease onset and vice versa. (9) Given the limited statistical power of our findings for post-event survival, our study supports the need for substantially larger well-phenotyped cohorts to differentiate effects of variants associated with CHD from post-event mortality.

An advantage of this study is that within the setting of the CHARGE Consortium we are able to evaluate and make comparisons between cross-sectional and prospective study designs, and to investigate all-cause mortality following cardiovascular events. There are differing, but overlapping, sample sizes across the various study designs: 27,349 participants from seven cohorts for prevalent outcomes, 55,736 participants from nine cohorts for incident outcomes, and 3,751 MI and CHD cases from six cohorts that contributed to the analysis of all-cause mortality. These differing sample sizes influence our power to detect associations, and inferences about similarities and differences across study designs could be due to biological differences or differences in sample sizes. This investigation of low-frequency and rare variants was limited to the variants included on the genotyping platforms (HumanHap300 Duo+ and HumanExome BeadChip, v1.0-1.2, Illumina, Inc., San Diego, CA) and was also limited to individuals of European ancestry. Additionally, although the variants on the genotyping platform and included in our gene-based tests were enriched for coding variants predicted to be causal, we cannot attribute causality to the variants or genes with novel associations. A strength of this study is that the quality of rare variant genotype calling was maximized by the joint clustering performed within CHARGE on thousands of samples (21).

In conclusion, this study comprehensively evaluated the relationship between autosomal genetic variation and prevalent and incident cardiovascular outcomes in participants of European ancestry in the context of the CHARGE consortium. We confirmed previously reported loci influencing heart disease risk as well as newly identified several loci associated with MI and CHD that warrant future investigation.

## Acknowledgments

The Age, Gene, Environment, Susceptibility Study (AGES) study has been funded by NIH contracts N01-AG-1-2100 and HHSN271201200022C, the NIA Intramural Research Program, Hjartavernd (the Icelandic Heart Association), and the Althingi (the Icelandic Parliament). The study is approved by the Icelandic National Bioethics Committee, VSN: 00-063. The researchers are indebted to the participants for their willingness to participate in the study.

The Atherosclerosis Risk in Communities study has been funded in whole or in part with Federal funds from the National Heart, Lung, and Blood Institute, National Institutes of Health, Department of Health and Human Services (contract numbers HHSN268201700001I, HHSN268201700002I, HHSN268201700003I, HHSN268201700004I and HHSN268201700005I). The authors thank the staff and participants of the ARIC study for their important contributions. Funding support for “Building on GWAS for NHLBI-diseases: the U.S. CHARGE consortium” was provided by the NIH through the American Recovery and Reinvestment Act of 2009 (ARRA) (5RC2HL102419).

Cardiovascular Health Study: This CHS research was supported by NHLBI contracts HHSN268201200036C, HHSN268200800007C, HHSN268201800001C, N01HC55222, N01HC85079, N01HC85080, N01HC85081, N01HC85082, N01HC85083, N01HC85086; and NHLBI grants U01HL080295, R01HL087652, R01HL105756, R01HL103612, R01HL120393, and U01HL130114 with additional contribution from the National Institute of Neurological Disorders and Stroke (NINDS). Additional support was provided through R01AG023629 from the National Institute on Aging (NIA). A full list of principal CHS investigators and institutions can be found at https://chs-nhlbi.org/.

The provision of genotyping data was supported in part by the National Center for Advancing Translational Sciences, CTSI grant UL1TR001881, and the National Institute of Diabetes and Digestive and Kidney Disease Diabetes Research Center (DRC) grant DK063491 to the Southern California Diabetes Endocrinology Research Center.

The Family Heart Study (FamHS) was supported by the grant R01-HL-117078 from the National Heart, Lung, and Blood Institute, and grant R01-DK-089256 from the National Institute of Diabetes and Digestive and Kidney Diseases.

The Framingham Heart Study (FHS) The National Heart, Lung and Blood Institute’s Framingham Heart Study is supported by contract N01-HC-25195.

GeneSTAR was supported by grants from the National Institutes of Health/National Heart, Lung and Blood Institute (HL49762, HL59684, HL071025, HL58625, U01 HL72518, HL089474, HL092165, HL099747, K23HL105897, K23HL094747, HL11006, and HL112064), National Institute of Nursing Research (NR0224103, NR008153), National Institute of Neurological Disorders and Stroke (NS062059), and by a grant from the National Center for Research Resources (M01-RR000052) to the Johns Hopkins General Clinical Research Center. Genotyping services were provided through the RS&G Service by the Northwest Genomics Center at the University of Washington, Department of Genome Sciences, under U.S. Federal Government contract number HHSN268201100037C from the National Heart, Lung, and Blood Institute.

MESA and the MESA SHARe project are conducted and supported by the National Heart, Lung, and Blood Institute (NHLBI) in collaboration with MESA investigators. Support for MESA is provided by contracts HHSN268201500003I, N01-HC-95159, N01-HC-95160, N01-HC-95161, N01-HC-95162, N01-HC-95163, N01-HC-95164, N01-HC-95165, N01-HC-95166, N01-HC-95167, N01-HC-95168, N01-HC-95169, UL1-TR-000040, UL1-TR-001079, UL1-TR-001420, UL1-TR-001881, and DK063491. Funding for SHARe genotyping was provided by NHLBI Contract N02-HL-64278. Provision of exome chip genotyping was provided in part by support of NHLBI contract N02-HL-64278 and UCLA CTSI UL1-TR001881, and the S.Calif DRC DK063491.

From the Rotterdam Study, the authors are grateful to the study participants, the staff, and the participating general practitioners and pharmacists. This work was supported by the Erasmus Medical Center and Erasmus University, Rotterdam; The Netherlands Organisation for the Health Research and Development (ZonMw); the Research Institute for Diseases in the Elderly (014-93-015, RIDE2); the Ministry of Education, Culture and Science; the Ministry for Health, Welfare and Sports; the European Commission (DG XII); the Municipality of Rotterdam; The Netherlands Organisation of Scientific Research (NWO) (175.010.2005.011, 911-03-012); the Netherlands Genomics Initiative (NGI) (NWO 050-060-810), the Netherlands Organisation for Scientific Research (NWO) (veni 916.12.154).

The Study of Health in Pomerania (SHIP) and SHIP-TREND both represent population-based studies. SHIP is supported by the German Federal Ministry of Education and Research (Bundesministerium für Bildung und Forschung (BMBF); grants 01ZZ9603, 01ZZ0103, and 01ZZ0403) and the German Research Foundation (Deutsche Forschungsgemeinschaft (DFG); grant GR 1912/5-1). SHIP and SHIP-TREND are part of the Community Medicine Research net (CMR) of the Ernst-Moritz-Arndt University Greifswald (EMAU) which is funded by the BMBF as well as the Ministry for Education, Science and Culture and the Ministry of Labor, Equal Opportunities, and Social Affairs of the Federal State of Mecklenburg-West Pomerania. The CMR encompasses several research projects that share data from SHIP. The EMAU is a member of the Center of Knowledge Interchange (CKI) program of the Siemens AG. SNP typing of SHIP and SHIP-TREND using the Illumina Infinium HumanExome BeadChip (version v1.0) was supported by the BMBF (grant 03Z1CN22). We thank all SHIP and SHIP-TREND participants and staff members as well as the genotyping staff involved in the generation of the SNP data. The Women’s Genome Health Study (WGHS) is supported by the National Heart, Lung, and Blood Institute (HL043851, HL080467, HL099355) and the National Cancer Institute (CA047988 and UM1CA182913), with collaborative scientific support and funding for genotyping provided by Amgen.

## Supporting information

**S1 Document. Characteristics of the participating cohorts.**

**S1 Table. Study participants’ characteristics for prevalent MI and CHD analysis.**

**S2 Table. Low-frequency and rare variants underlying top signals from gene-based analysis for prevalent events.**

**S3 Table. Study participants’ characteristics for incident MI and CHD analysis.**

**S4 Table. Low-frequency and rare variants underlying top signals from gene-based analysis for incident events.**

**S5 Table. Study participants’ characteristics for post MI and CHD mortality analysis.**

**S6 Table. Prevalent and incident findings in relation to corresponding outcomes.**

